# Branching Suppresses Self-Aggregation while Preserving Membrane Association in SET-M33: An Atomistic Molecular Dynamics Study

**DOI:** 10.64898/2026.10.12.757849

**Authors:** Saha Satvati, Marta Rigoli, Sara Fortuna, Chiara Falciani, Alessandro Pini, Walter Rocchia

## Abstract

Branched antimicrobial peptides offer a promising strategy for improving the stability and activity of membrane-active peptides, but how branching alters self-association and membrane binding remains unclear. We used all-atom molecular dynamics simulations to compare tetra-branched SET-M33 with its linear branch sequence Q33 (KKIRVRLSA) in water and at an anionic bacterial-membrane model, to do this, systems containing equivalent numbers of Q33 sequence units were simulated. Q33 formed dimers and higher-order oligomers, particularly at the membrane interface and under hydration-tuned enhanced sampling. In contrast, SET-M33 remained more dispersed, as the branched scaffold and its high cationic charge limited peptide-peptide association while allowing multivalent membrane binding. Both architectures associated rapidly with the bilayer but remained predominantly surface-bound, without persistent pore formation or complete insertion. Q33 showed stronger membrane-induced helical stabilization, whereas SET-M33 retained a more heterogeneous conformational ensemble. Overall, branching primarily altered lateral peptide organization while preserving close membrane association, suppressing uncontrolled self-aggregation and providing an atomistic rationale for the distinct behavior of SET-M33 relative to Q33.

## INTRODUCTION

Antimicrobial peptides (AMPs) are being actively investigated as alternatives or complements to conventional antibiotics in response to the global rise of multidrug-resistant bacterial infections. More than 5000 natural, synthetic, and computationally predicted AMPs have been catalogued in the Antimicrobial Peptide Database, reflecting the rapid expansion of this field and the continuing discovery of new peptide scaffolds ^1–4^. Despite their promise, translation of AMPs into clinical use remains limited by toxicity, proteolytic instability, short half-life, and production cost ^5^. These limitations have motivated the design of peptidomimetic and multivalent architectures, including dendrimeric multiple-antigen peptides (MAPs), which can improve biological stability and potency relative to the corresponding monomeric sequences ^6^.

The improved stability of branched MAPs is commonly attributed to their multivalent architecture, in which several copies of a bioactive sequence are displayed from a central core ^7^. This organization can increase the local effective concentration of membrane-active motifs, extend biological half-life, and enhance bactericidal activity ^8^. For cationic AMPs, the earliest stage of bacterial targeting is generally driven by electrostatic attraction to negatively charged membrane components. Membranes enriched in anionic lipids interact more strongly with cationic amphiphilic peptides, which can promote surface accumulation, peptide aggregation, and membrane perturbation ^9–12^.

The balance between cationic charge and hydrophobicity is central to AMP function. Lysine, arginine, and histidine residues contribute to electrostatic recognition of bacterial surfaces, whereas hydrophobic residues support interfacial partitioning and bilayer penetration ^13^. Gram-positive bacteria contain anionic teichoic acids in the cell wall, whereas Gram-negative bacteria expose lipopolysaccharides (LPS) in the outer leaflet of the outer membrane; in both cases, cytoplasmic membranes contain anionic phospholipids such as phosphatidylglycerol, phosphatidylserine, and cardiolipin ^14–16^. Increasing the net positive charge of a peptide often enhances antibacterial activity, but residue patterning and amphipathic organization are equally important determinants of selective membrane binding and disruption ^17–20^. Proposed mechanisms include pore formation, carpet-like surface disruption, membrane thinning, charged-lipid clustering, targeting of oxidized lipids, and anion-carrier-like behavior ^21, 22^.

SET-M33 (hereafter M33) is a tetra-branched antimicrobial peptide originally identified through phage-display screening ^23^. M33 is active against several Gram-negative pathogens, including *Pseudomonas aeruginosa*, *Klebsiella pneumoniae*, and *Acinetobacter baumannii* ^24^. Experimental studies indicate that M33 binds strongly to LPS in the outer membrane of *P. aeruginosa* and that peptide accumulation on the membrane surface is associated with cytoplasmic damage and bacterial death ^25^. M33 consists of a lysine-based core carrying four copies of the nine-residue sequence Q33 (KKIRVRLSA) ^26^. Compared with the monomeric Q33 peptide, the branched construct shows increased resistance to proteases and peptidases while maintaining antibacterial activity ^27^.

Experimental characterization of M33 at atomic resolution is challenging because highly cationic, membrane-active peptides are difficult to study under conditions that avoid aggregation, precipitation, or membrane remodeling. Molecular dynamics (MD) simulations can complement experiments by resolving peptide conformational dynamics, peptide-peptide contacts, and peptide-membrane interactions at the atomic level ^28–30^. However, the relationship between branched topology, aggregation propensity, and membrane binding has not yet been fully clarified for M33-like constructs.

In this work, we used all-atom MD simulations to compare the behavior of linear Q33 and tetra-branched M33 in water and at an anionic model membrane. To better understand the effect of branching, systems were constructed with equivalent numbers of Q33 sequence units arranged either as free linear peptides or as branches of M33. We analyzed peptide aggregation, membrane coverage, secondary-structure evolution, peptide-membrane distance distributions, and residue-level hydration. The results indicate that linear Q33 readily forms membrane-associated dimers and higher-order oligomers, whereas the branched M33 architecture suppresses uncontrolled aggregation while preserving strong membrane association. These findings provide a molecular rationale for the distinct behavior of linear and branched antimicrobial peptide constructs.

## COMPUTATIONAL METHODOLOGY

### Peptide Models and System Construction

The linear peptide Q33 was modeled with the sequence KKIRVRLSA.

The branched peptide M33 consists of four identical Q33 (branches) covalently linked through a three-lysine branching core represented as (KKIRVRLSA)4K2K-betaA-O-. The lysine residues forming this core exhibit noncanonical connectivity because both backbone and side-chain amino groups participate in the branching amide linkages. Consequently, the standard lysine residue definitions available in conventional AMBER force-field libraries could not be directly applied to the branching residues. Standard force-field parameters were retained for the conventional amino acid residues of the peptide branches, whereas a dedicated force-field library was generated for the nonstandard lysine core using R.E.D. Server Development/PyRED ^31–34^.

The branched core was parameterized using the molecular-fragment approach implemented in PyRED, in which a complex molecular structure is decomposed into elementary building blocks representative of its local chemical environment. Cleavage sites corresponding to peptide bonds were terminated with acetyl (CH₃CO–) and N-methylamide (–CONHCH₃) capping groups. This strategy preserves the local amide environment of each fragment while avoiding quantum-mechanical optimization of the complete, highly branched peptide as a single molecular entity. Appropriate charge constraints were imposed on the capping groups during electrostatic-potential fitting, and the atoms belonging to these groups were subsequently removed from the resulting force-field fragments. The parameterized fragments were then connected according to the native branching topology of M33 to reconstruct the three-lysine core.

The goal of the model design was to compare the same bioactive sequence displayed either as independent linear peptides or as covalently linked branches of a single scaffold.

Atomic partial charges were obtained by fitting the quantum-mechanically calculated molecular electrostatic potential using the restrained electrostatic potential (RESP) procedure. For calculations performed with the standard RESP-A1 protocol implemented in PyRED, molecular geometries were optimized and electrostatic potentials were calculated at the HF/6-31G(d) level, followed by a two-stage RESP fit using a Connolly molecular surface. The resulting charges and residue definitions were incorporated into an AMBER-compatible force-field library and combined with the standard parameters used for the peptide branches.

Because electrostatic-potential-derived charges can depend on molecular conformation, the starting geometries of the elementary building blocks were selected to minimize artificial contacts between groups that are spatially separated in the assembled peptide. Following reconstruction of the complete branched peptide, the four peptide arms were initially oriented away from one another to avoid introducing preformed interbranch interactions into the starting structure. This arrangement was used solely as an unbiased initial configuration; conformational relaxation of the peptide was subsequently allowed during the molecular dynamics equilibration and production simulations. The Lysines in each branch of SET-M33 as well as these in Q33 were assumed to be in the protonated state. The N-termini of the branches and all the termini of Q33 peptides were in the charged state. The overall net charge of M33 was +19, and that of Q33 was +4.

An anionic bilayer containing 245 phospholipids per leaflet was constructed with a lipid ratio of 2:1.3:1 for 1,2-dioleoyl-sn-glycero-3-phosphoethanolamine (DOPE), 1,2-dipalmitoyl-sn-glycero-3-phosphoglycerol (DPPG), and 1,2-dipalmitoyl-sn-glycero-3-phosphoethanolamine (DPPE), respectively (see Figure 1). This corresponds to an anionic bacterial-membrane model inspired by reported P. aeruginosa lipid compositions. 35. Initial membrane coordinates were generated with PACKMOL-Memgen ^36^, and Packmol and tLEaP ^37–39^ were used for system assembly and topology preparation.

**Figure 1.**
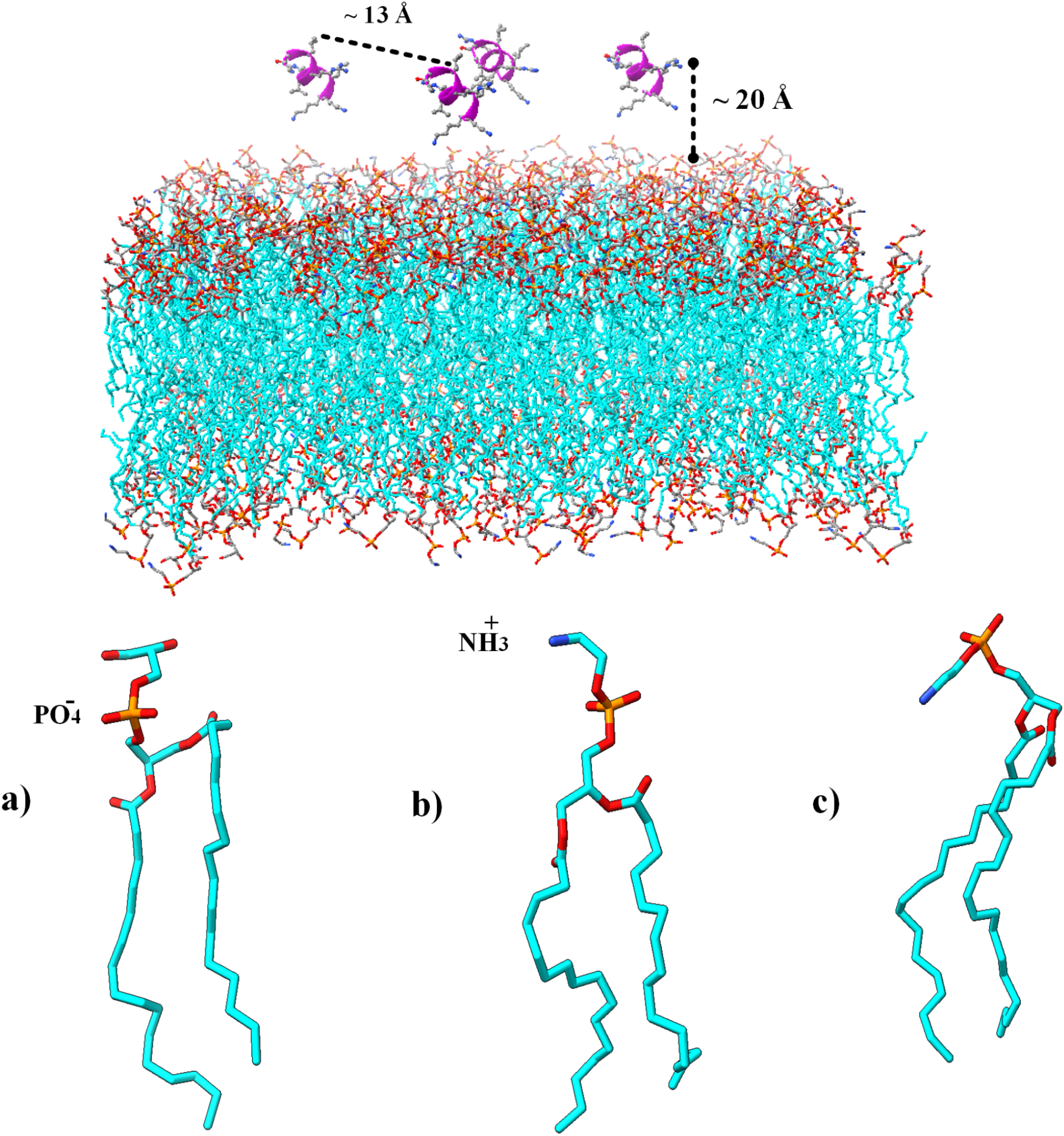
Initial configuration of representative peptide–membrane systems. The membrane contains DOPE, DPPG, and DPPE lipids in a 2:1.3:1 ratio. Peptides were initially placed in the aqueous phase at least 20 Å from the membrane surface with randomized orientations. Water and ions are omitted for clarity. Lipid structures are shown for (a) DPPG, (b) DPPE, and (c) DOPE.

Peptides were placed in water above the outer side of the membrane at least 20 Å from the membrane surface and with randomized orientations unless otherwise specified. Each peptide-membrane condition was simulated in three independent replicas. Systems were solvated with TIP3P water and 150 mM KCl to neutralize the system and mimic the ionic conditions used experimentally. The full list of simulated systems is provided in Table 1.

**Table 1.**
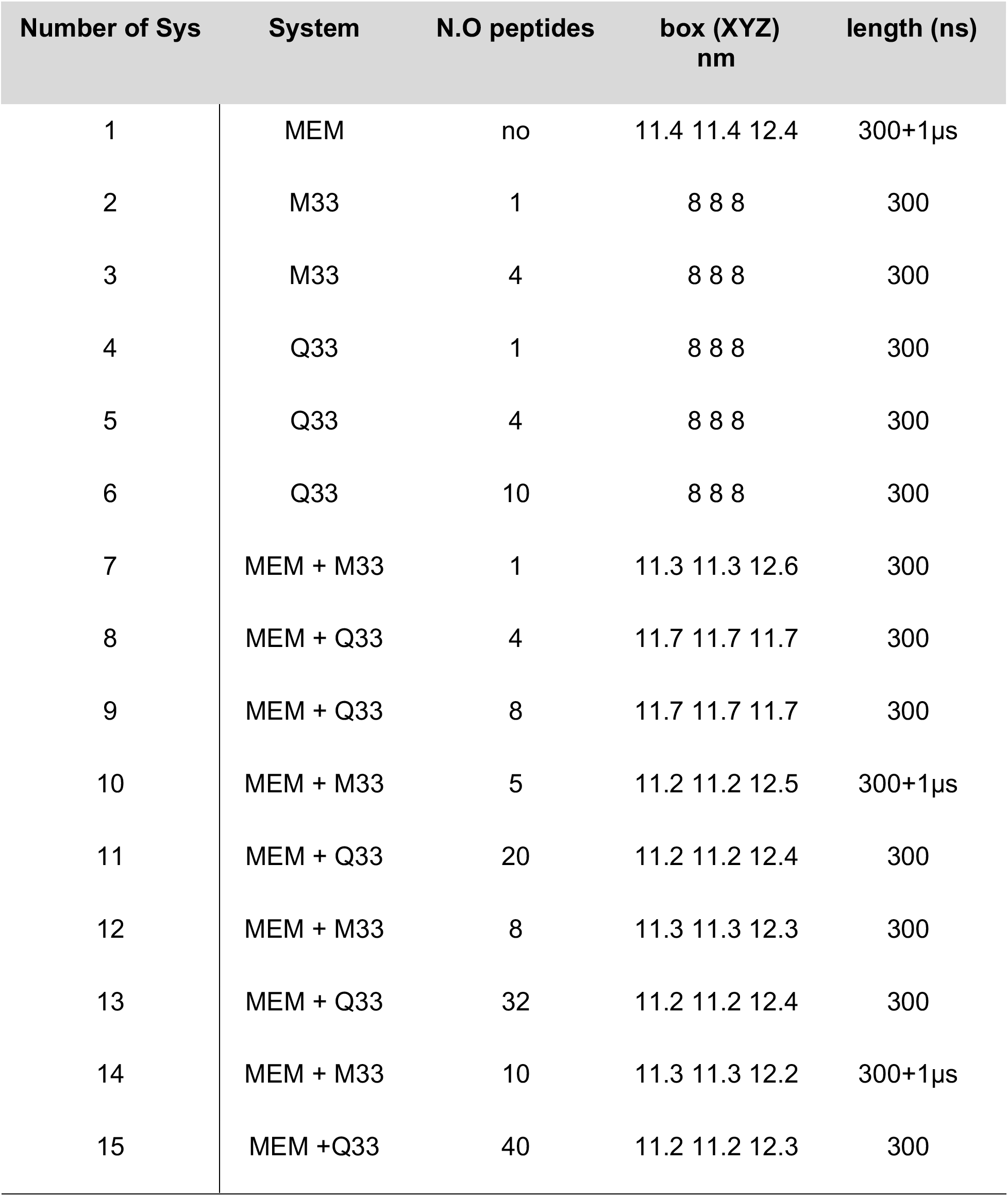
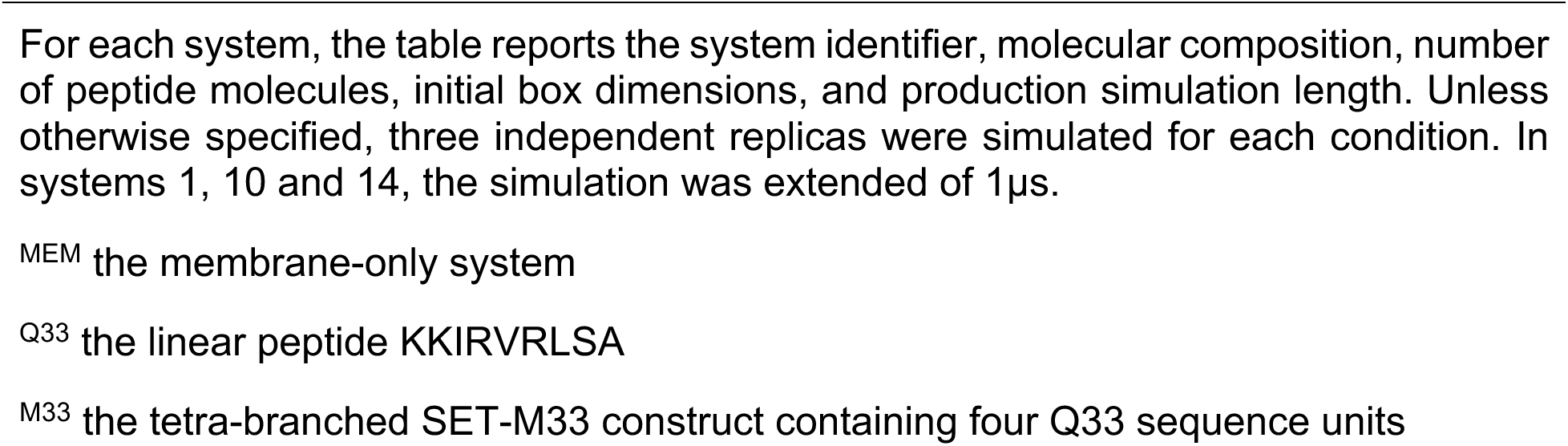
Simulated systems used for all-atom molecular dynamics simulations.

### Molecular Dynamics Protocol

All simulations were performed with GROMACS 2024.3 ^40–44^. Peptides were described with the AMBER ff14SB force field, and lipids were described with Lipid21 parameters ^45, 46^. Energy minimization was carried out with the steepest-descent algorithm. The systems were then equilibrated in three NVT stages, during which heavy atoms of the peptides and lipids were position restrained while the temperature was increased from 100 to 300 K with the Nose-Hoover thermostat ^47^.

Subsequent NPT equilibration was performed at 1 bar using the semi-isotropic Parrinello-Rahman barostat with a pressure-coupling time constant of 5 ps ^48^. During the first stage, all heavy atoms of peptide were restrained for 10 ns, then all position restraints were removed before the production simulation. A 2 fs integration time step was used, and bonds involving hydrogen atoms were constrained with the LINCS algorithm ^49^. Production simulations were performed in the NPT ensemble at 300 K and 1 bar. Unless otherwise stated, each production trajectory was 300 ns long; selected high-density M33 systems were extended to 1 us to assess slow peptide-peptide association events. Electrostatic and van der Waals interactions were truncated at 1.2 nm, and long-range electrostatics were treated with the smooth particle-mesh Ewald method using a grid spacing of 0.16 nm.

### Hydration-Tuning Enhanced Sampling

To probe the aggregation propensity of Q33, hydration-tuning enhanced sampling was applied to the system containing 10 Q33 peptides in water. The method introduces a screened pairwise electrostatic-like bias between predefined solute groups, for details see reference ^50, 51^, allowing the effective hydration of selected species to be tuned and thereby accelerating association events. For the present application, the two groups were the heavy atoms of the peptides, and the Oxygen atoms of the surrounding water molecules, respectively. By recording the level of reduction of peptide-water interaction needed to observe aggregation, we could infer the relative aggregation propensity in a qualitative and accelerated fashion. The reduction is calculated with respect to a reference taken as average hydration value calculated over the 50 ns unbiased MD simulation. Related biasing approaches have been used to accelerate protein-ligand association and binding ^51^.

For the 10-Q33 system, three biased trajectories were performed at 90%, 80%, and 50% of the reference mean hydration obtained from the corresponding unbiased simulation. To enable comparison with the branched architecture, the same protocol was applied to the 4-M33 system.

Because hydration tuning modifies the effective solvation environment, the resulting trajectories were not used to infer equilibrium populations. Instead, they were used to identify possible association pathways and interaction motifs that can be compared with those sampled in unbiased solution and membrane simulations.

### Trajectory Analysis

Trajectory analyses were performed with GROMACS and VMD 1.9.4 52. Before analysis, trajectories were corrected for periodic boundary conditions, molecules were made whole, and membrane-containing systems were centered with respect to the bilayer. Unless otherwise specified, analyses were performed over the equilibrated portion of each production trajectory, and reported distributions were averaged over the three independent replicas. Peptide-peptide contacts were quantified using heavy-atom contact maps generated with gmx mdmat, applying a distance cutoff of 0.4 nm. Contacts were calculated between residues belonging to different peptide molecules or, in the case of M33, between different branched peptide constructs. This analysis was used to identify residue pairs contributing to peptide-peptide association in solution and at the membrane interface.

Lateral peptide organization on the membrane was characterized using center-of-mass-based radial distribution functions (RDFs) and cumulative coordination numbers (CNs). For each system, one center of mass was assigned to each peptide molecule included in the analysis. RDFs were calculated in the membrane plane using the gmx rdf –xy option, considering only peptides associated with the same leaflet. Architecture-specific distance cutoffs were used to define close lateral association and first-neighbor shells. For M33, cutoffs of 3.2 and 4.0 nm were used, reflecting the larger lateral extension of the tetra-branched construct. For Q33, cutoffs of 1.7 and 2.0 nm were used, reflecting the smaller size of the linear peptide. These cutoffs were selected from peptide dimensions and the corresponding RDF profiles and were used to compare local peptide crowding within each architecture.

Peptide-membrane association was quantified from the minimum distance between peptide heavy atoms and phosphate atoms of the upper membrane leaflet using gmx distance and gmx mindist. A minimum-distance threshold of 0.4 nm was used to define close peptide-membrane contact. This metric was interpreted as contact persistence or bound-state occupancy not as evidence of irreversible binding.

Membrane coverage was estimated from the solvent-accessible surface area (SASA) of the lipid bilayer using the double-cubic-lattice method implemented in gmx sasa. Because peptide adsorption shields lipid headgroups from water, lower membrane SASA values were interpreted as greater peptide-induced membrane coverage. Peptide compactness and spatial spreading were further assessed using radius-of-gyration analysis where appropriate. Secondary-structure evolution was analyzed with gmx dssp, using the DSSP assignment of helix, β-structure, turn, bend, and coil states along the trajectory. For M33, secondary-structure content was evaluated separately for the four Q33 branches and then averaged where needed to enable comparison with linear Q33 systems. Residue-level hydration was quantified by counting water molecules within 0.4 nm of each residue using gmx select and in-house Python scripts. Reduced hydration was interpreted as increased interfacial association or partial burial, whereas elevated hydration indicated greater solvent exposure. For membrane-associated systems, hydration profiles were compared with residue identity and peptide orientation to identify hydrophobic residues preferentially positioned near the lipid interface and charged residues remaining exposed to water and lipid headgroups.

For scalar observables and probability distributions, uncertainty was estimated from variation among the three independent replicas, and replica-averaged values are reported unless otherwise specified.

## RESULTS and DISCUSSION

### Q33 Aggregation in Solution and at the Membrane

Previous experimental work by Falciani and co-workers showed that M33 and Q33 differ markedly in stability and antibacterial performance against Gram-negative bacteria ^26^. We therefore first examined the intrinsic behavior of the linear Q33 peptide in solution and then evaluated how peptide loading affects Q33 association at the membrane interface.

In unbiased solution simulations, the 10-Q33 system did not undergo large-scale aggregation within the sampled timescale, indicating that formation of compact oligomers is slow under fully hydrated conditions. Hydration-tuned enhanced sampling was therefore used as an accelerated probe of aggregation-prone configurations rather than as a direct measure of equilibrium aggregation kinetics. At milder hydration reductions of 90% and 80% transient peptide-peptide encounters is increased, leading to the formation of two dimers. However, no persistent large aggregates were observed in either system. Under the strongest dehydration condition, corresponding to 50% of the reference hydration level, Q33 rapidly formed compact assemblies containing up to nine peptides. The mass-density maps showed a more localized peptide distribution in the biased trajectory than in the unbiased equilibrated ensemble, consistent with reduced spatial fluctuations and enhanced clustering (Figure 2).

**Figure 2.**
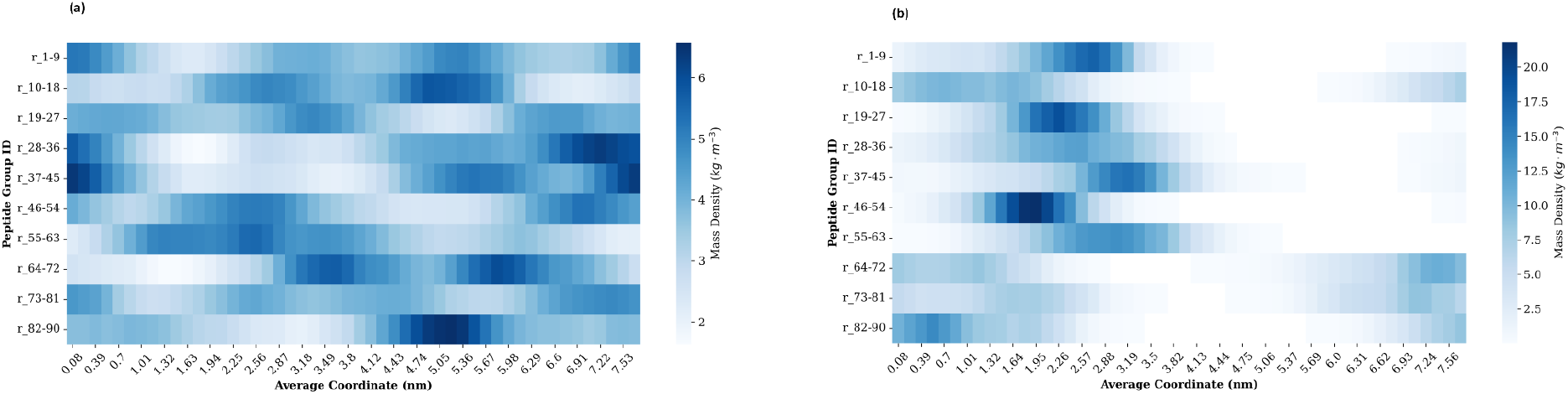
Peptide mass-density maps for the 10-Q33 system in water. (a) Mass-density distribution obtained from the unbiased equilibrated simulation. (b) Mass-density distribution obtained under hydration-tuned enhanced sampling at 50% of the reference hydration level. The biased trajectory promotes a more localized peptide distribution, consistent with compact aggregate formation.

Residue-level contact analysis of the aggregated states identified recurrent interactions involving hydrophobic residues, including alanine, isoleucine, and leucine, together with polar backbone contacts. In several peptide-peptide interfaces, backbone carbonyl and amide groups from residues carrying lysine or arginine side chains contributed to close intermolecular approaches. During aggregation, center-of-mass distances between selected peptide pairs decreased below approximately 2 nm, while minimum heavy-atom distances stabilized near or below 0.5 nm, confirming formation of direct intermolecular contacts (Figure 3). These biased trajectories therefore indicate which contact motifs can stabilize Q33 assemblies when peptide desolvation and close approach are promoted.

**Figure 3.**
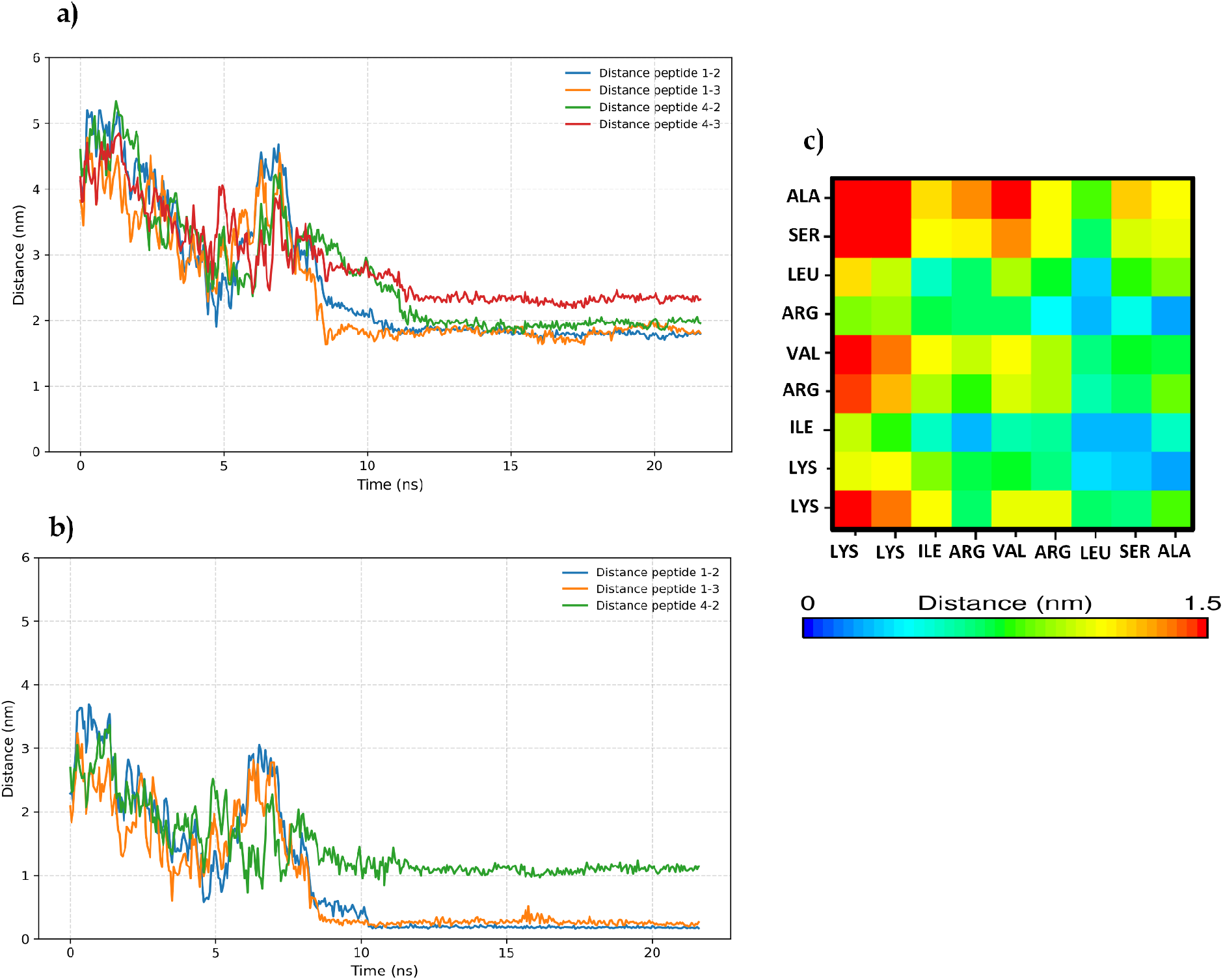
Peptide–peptide contacts in the hydration-tuned 10-Q33 simulation. (a) Center-of-mass distances between a reference Q33 peptide and selected neighboring peptides. (b) Minimum heavy-atom distances between representative peptide pairs. (c) Residue–residue contact map for a representative Q33 dimer. Contacts were calculated using a 0.4 nm heavy-atom cutoff.

Q33 association was also observed in membrane-containing simulations, where peptide adsorption onto the anionic bilayer favored local peptide crowding. In the 4-Q33 membrane system, three peptides formed an oligomeric assembly early during the production trajectory and remained associated after establishing membrane contact (Figure 4). In the 8-Q33 system, dimers and occasional tetramers were observed at the membrane interface (Figure 5). At higher peptide numbers, namely 20, 32, and 40 Q33 peptides, aggregation became more pronounced, and produced dense peptide-rich regions on the membrane surface (Figure 6). These clusters were not uniformly distributed over the bilayer, indicating that increased Q33 loading promotes aggregation-driven adsorption rather than homogeneous membrane coverage.

**Figure 4.**
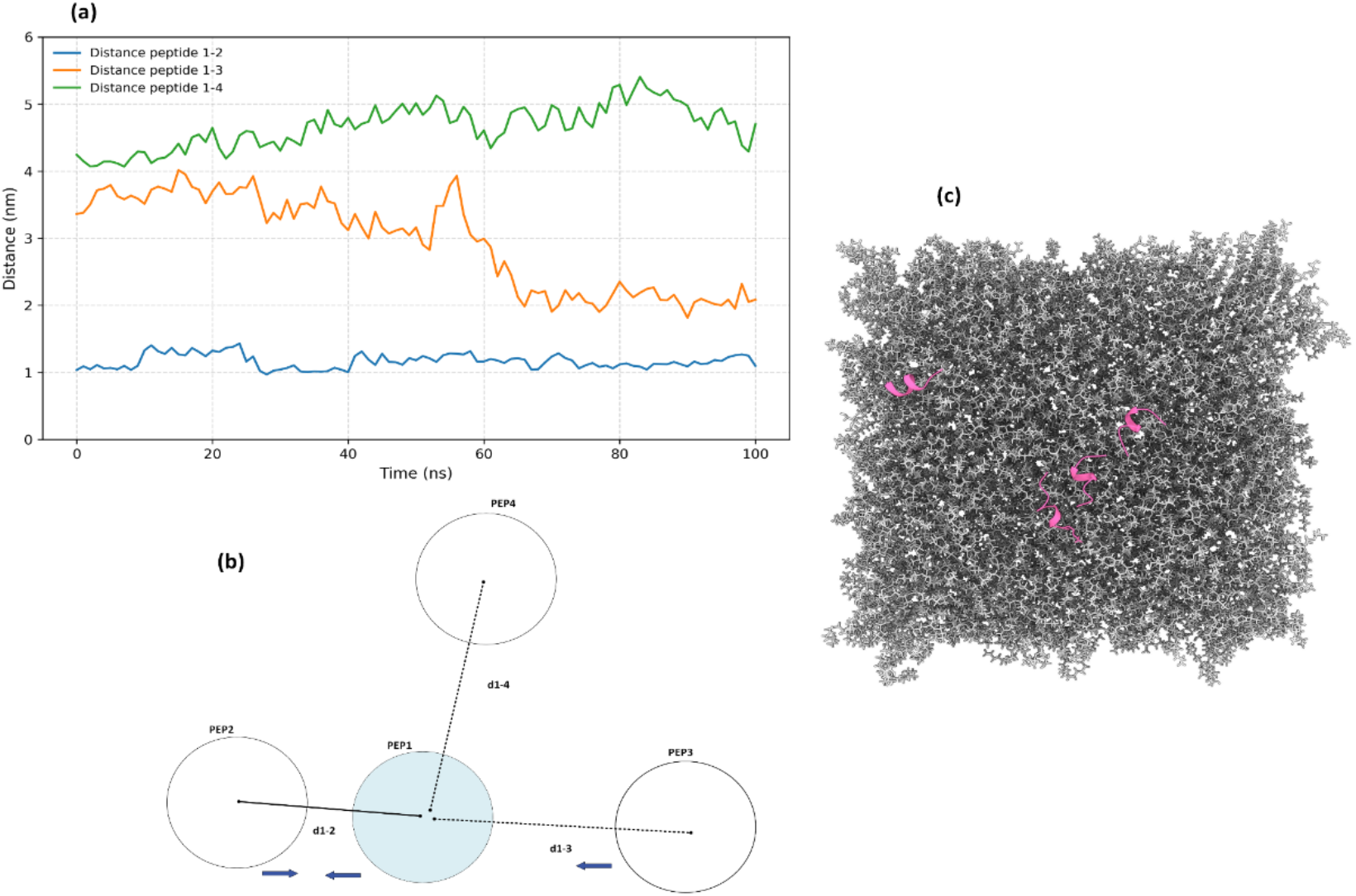
Four Q33 peptides interacting with the membrane. (a) Center-of-mass (COM) distances between a reference peptide (Pep1) and the other three peptides (Pep 2, 3, 4). (b) Relative positions of three peptides with respect to the reference peptide; blue arrows indicate interacting peptide pairs. (c) Representative snapshot of the peptides on the membrane, colored according to their secondary structure.

**Figure 5.**
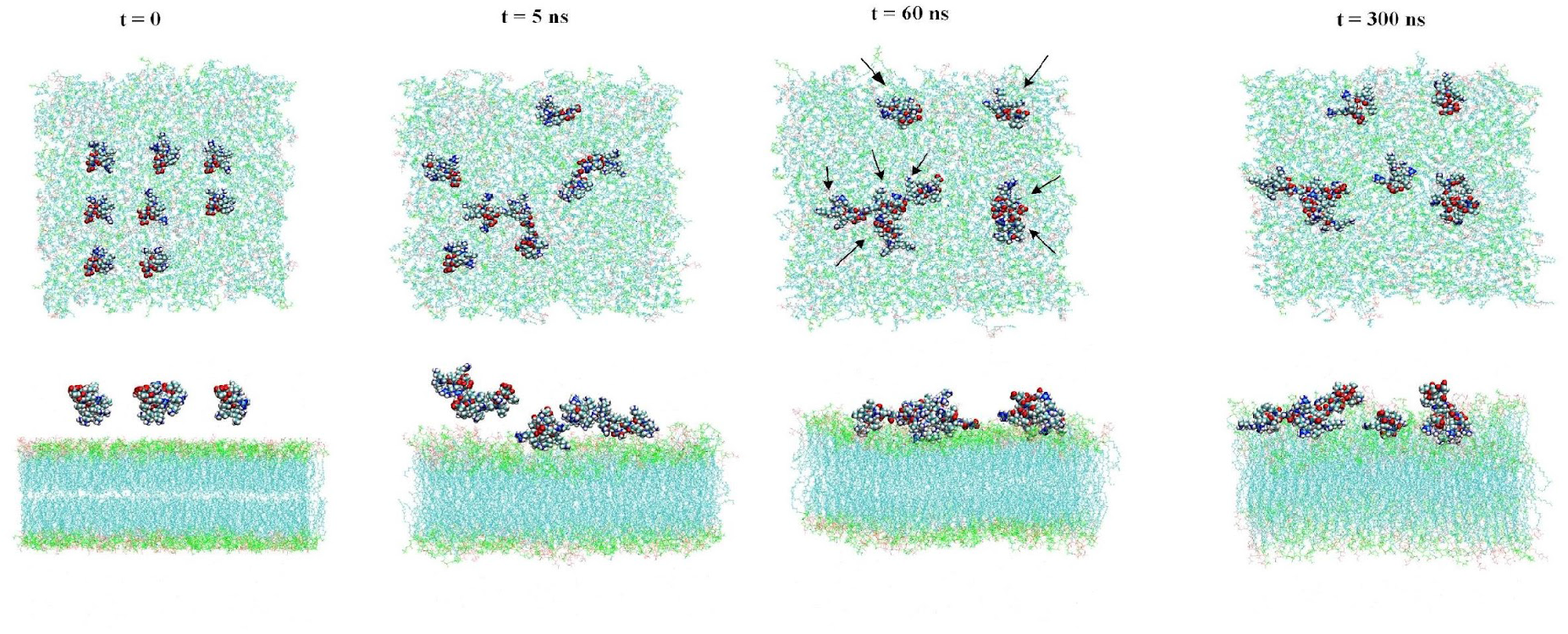
Representative snapshots of the 8-Q33-membrane system at 0, 5, 60, and 300 ns following removal of positional restraints. Eight Q33 peptides were initially placed above the membrane. As the simulation expanded, peptide aggregation occurred on the bilayer surface, with dimers and tetramers generated within 60 ns.

**Figure 6.**
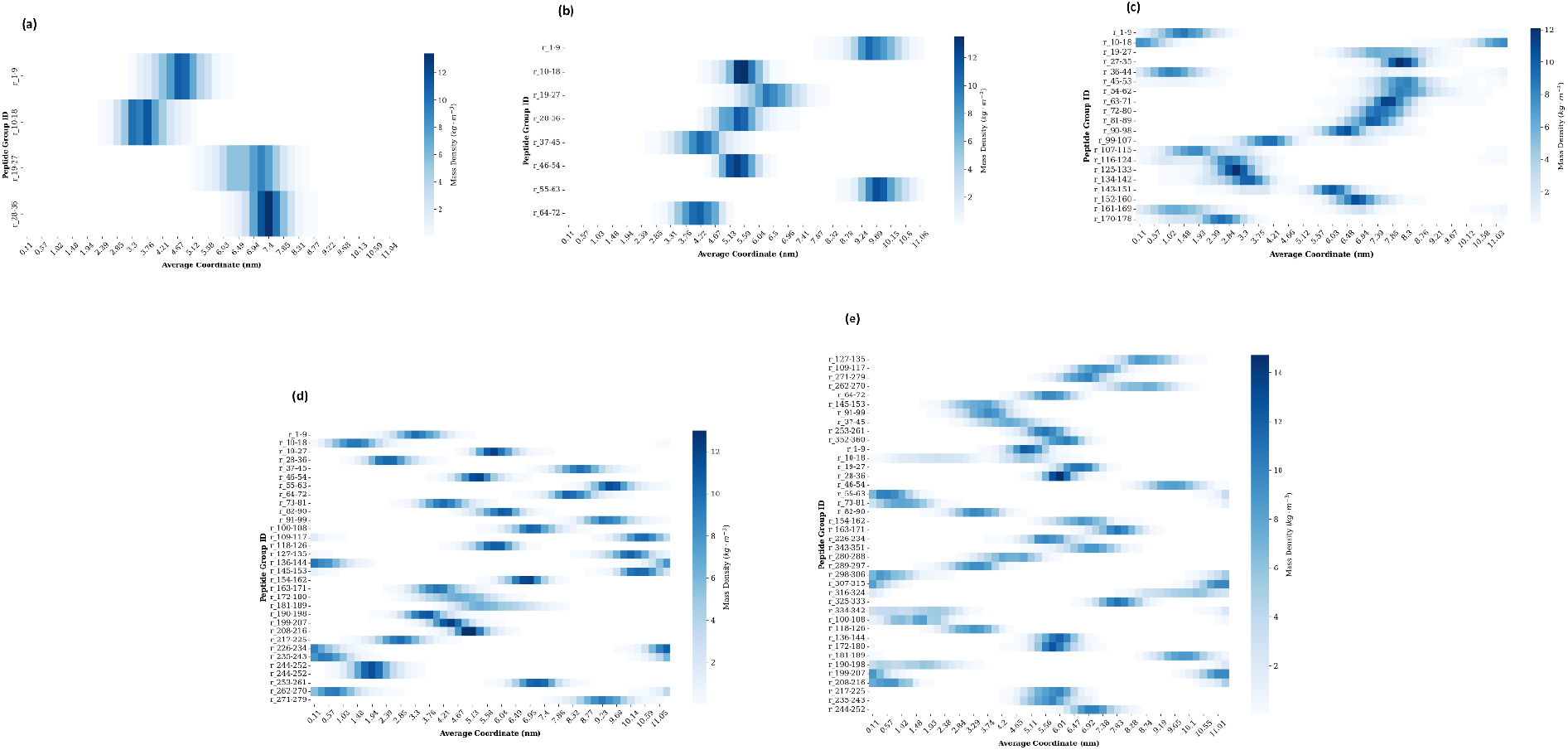
Lateral density maps of Q33 peptides on the membrane surface. Time-averaged peptide density distributions are shown for systems containing (a) 4, (b) 8, (c) 20, (d) 32, and (e) 40 Q33 peptides. Higher-density regions correspond to preferential peptide accumulation and peptide-rich clusters at the membrane interface.

Contact maps from membrane-associated Q33 assemblies were consistent with the motifs identified in solution (Figure 7). Dimeric contacts involved hydrophobic residues together with interactions involving lysine– and arginine-containing segments. At higher local peptide densities, charged residues appeared to contribute to the recruitment of additional peptides, allowing dimers to act as nucleation sites for trimers, tetramers, and larger surface-associated assemblies. Overall, the Q33 simulations show that the linear architecture permits close peptide-peptide packing both before and during membrane binding. This behavior contrasts with the more dispersed membrane association observed for the branched M33 scaffold, discussed below.

**Figure 7.**
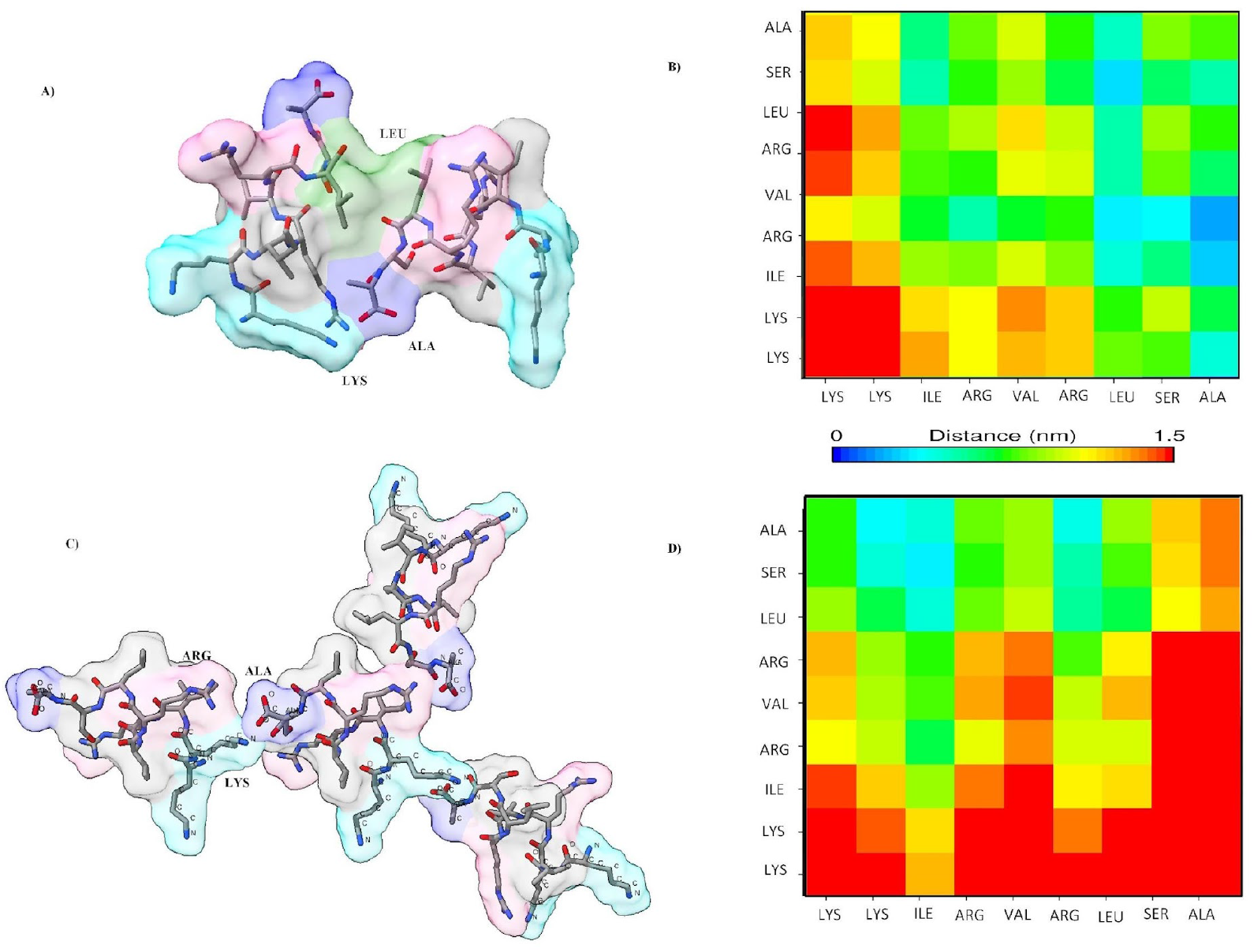
Progression of Q33 peptide aggregation from solution to membrane association in the system containing eight Q33 peptides and an anionic membrane. (A) Representative initial interaction between two Q33 peptides in solution, prior to membrane association. Residues involved in the peptide-peptide interface are highlighted, including alanine (blue), lysine (cyan), and arginine (pink). (B) Residue–residue contact map of the two selected Q33 peptides forming a stable dimeric assembly. (C) Representative snapshot showing the persistence and further association of Q33 peptides after membrane binding, resulting in a four-peptide assembly at the membrane surface. (D) Contact map of two Q33 within the four-Q33 assembly, highlighting close contacts involving alanine and positively charged lysine and arginine residues that contribute to stabilization of the aggregate.

### M33 Membrane Association and Suppressed Self-Aggregation

The behavior of tetra-branched M33 differed substantially from that of linear Q33. In aqueous solution, M33 molecules remained largely dispersed and did not form persistent higher-order aggregates over the sampled timescales. This reduced self-association is consistent with the high density of cationic lysine and arginine residues displayed across the branches, which likely disfavors close M33-M33 approach, together with the steric and topological constraints imposed by the branched scaffold. Increasing the number of M33 molecules in solution did not produce the extensive clustering observed for Q33, and the density maps supported a comparatively dispersed peptide population (Figure 8a).

**Figure 8.**
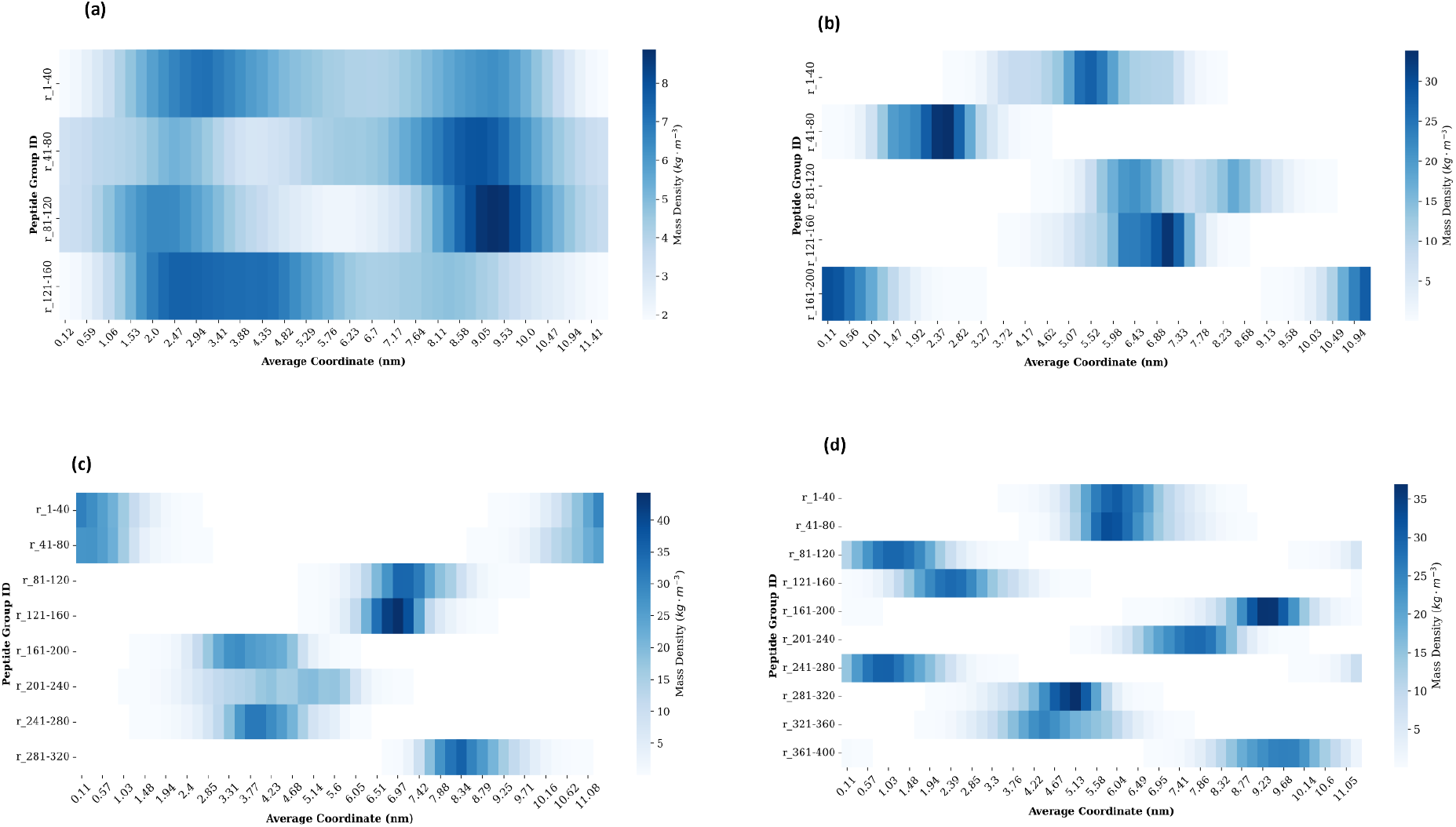
Spatial distribution of M33 peptides in solution and membrane systems. Density maps are shown for (a) 4 M33 peptides in water, (b) 5 M33 peptides with membrane, (c) 8 M33 peptides with membrane, and (d) 10 M33 peptides with membrane. Compared with Q33, M33 displays a more dispersed spatial distribution, particularly at low and intermediate peptide loading.

In membrane-containing systems, M33 associated rapidly with the anionic bilayer after release of positional restraints. Membrane binding reduced lateral diffusion but did not lead to extensive M33-M33 oligomerization at low and intermediate peptide loadings (Figure 8b,c). In contrast to Q33, whose membrane association was accompanied by frequent peptide-peptide clustering, M33 binding was dominated by multivalent interactions between its cationic branches and the anionic lipid headgroups. The branched architecture allows multiple Q33 segments to contact the membrane simultaneously, while limiting the exposure of hydrophobic and core regions to neighboring M33 molecules.

At higher M33 concentrations, M33-M33 contacts became more frequent but remained strongly dependent on molecular orientation. Contact-map analysis of the 10-M33 membrane system showed that the most persistent contacts involved favorable exposure of hydrophobic residues, particularly isoleucine and leucine, and/or interactions involving the central lysine-based core (Figure 9). In the extended 1 μs simulations, occasional pairwise associations were observed under specific configurations, but these contacts did not develop into the extensive oligomeric assemblies observed for Q33. Thus, M33 crowding appears to require more restrictive geometrical conditions and occurs less readily than aggregation of the linear peptide.

**Figure 9.**
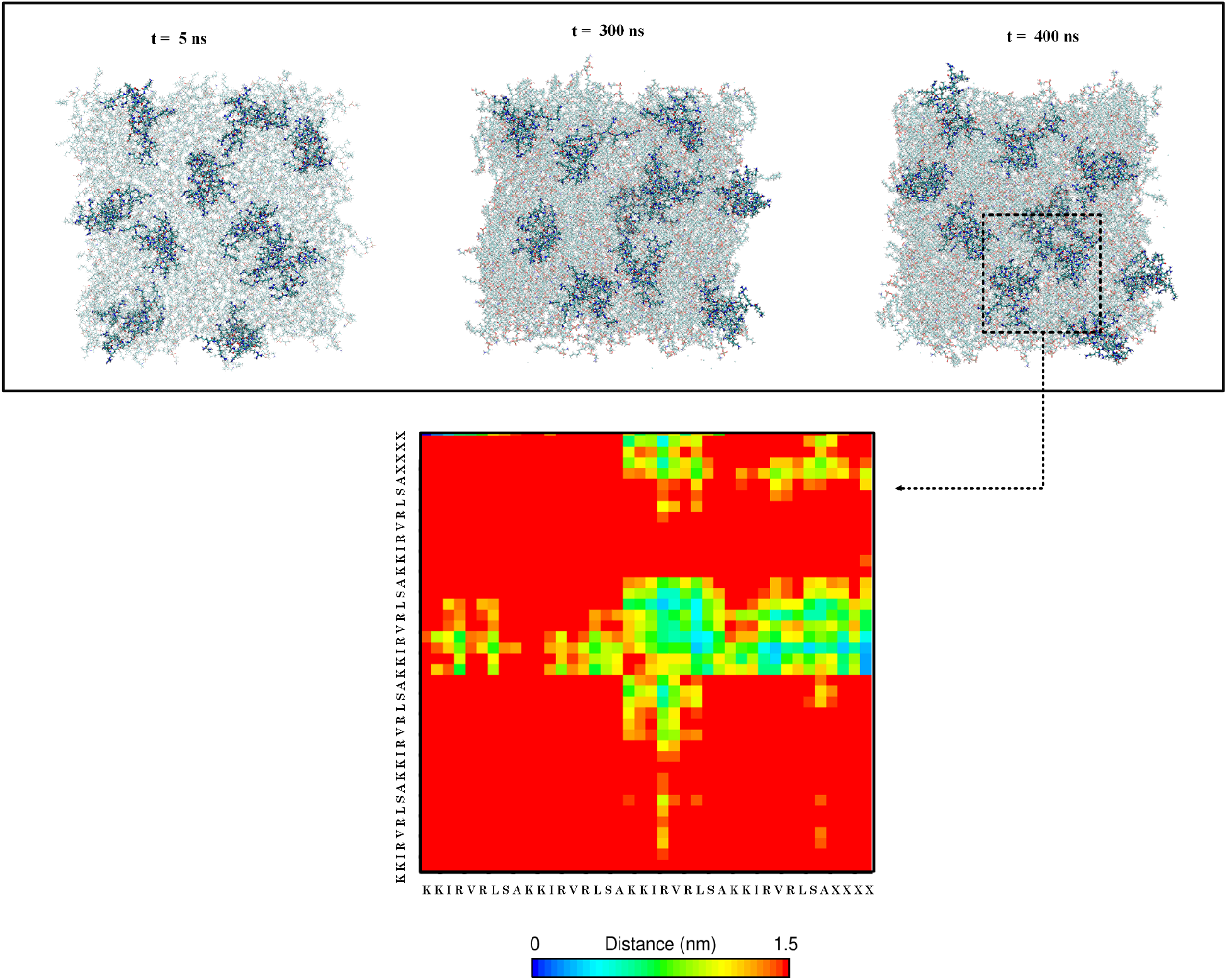
Representative configurations and contact analysis of the 10-M33 membrane system. Snapshots show selected M33 molecules in the membrane-associated state during the extended trajectory. Pairwise contacts were rare and orientation-dependent, involving hydrophobic residues and/or the central core when favorable molecular arrangements were sampled. No extensive higher-order M33 oligomerization comparable to Q33 clustering was observed.

This interpretation is supported by the lateral RDF and cumulative coordination number analyses, which show only moderate increases in M33 first-neighbor coordination with peptide loading, whereas Q33 displays stronger short-range enrichment and progressively higher local coordination.

Overall, two classes of M33-M33 interactions were identified. The first consisted of transient contacts involving charged residues, which were limited by electrostatic repulsion and rapid reorientation. The second involved more persistent contacts between hydrophobic residues and/or the core region when favorable orientations exposed these groups to another M33 molecule. These observations indicate that branching changes the dominant interaction pathway: whereas Q33 competes between peptide-peptide aggregation and peptide-membrane binding, M33 preferentially engages the membrane while suppressing uncontrolled self-association.

In contrast to the Q33 case, forcing a reduction in hydration in the M33 system did not produce detectable peptide-peptide associations within the 50 ns simulation time. Thus, under the above described conditions, the bias had no measurable effect on oligomerization of the branched peptide. This behavior is consistent with steric/topological constraints and slower intermolecular encounter dynamics. At milder hydration reduction levels of 90% and 80%, Q33 showed increased peptide-peptide encounters relative to the unbiased trajectory but did not form the compact nine-peptide aggregate observed at 50% hydration within the same simulation time. Thus, the 50% condition was used to characterize the residue-level interaction motifs associated with the strongly aggregated state, whereas the milder conditions served as controls confirming the progressive nature of the hydration-driven association response.

### Membrane Coverage, Lateral Organization, and Contact Distance

To determine how peptide architecture affects membrane engagement beyond simple peptide adsorption, we analyzed three complementary observables: membrane solvent-accessible surface area (SASA), lateral peptide organization on the membrane, and minimum peptide–membrane distance. Together, these analyses distinguish membrane shielding, surface crowding, and contact distance.

Membrane coverage was first evaluated from the SASA of the lipid bilayer in the presence of Q33 and M33 peptides (Figure 11a–d). Because adsorbed peptides shield lipid headgroups from water, lower membrane SASA values indicate greater peptide-induced surface coverage. At the lowest peptide loading, the SASA distributions for 4 Q33 and 1 M33 overlapped substantially, suggesting comparable membrane shielding when the same number of Q33 sequence units was present. At intermediate loading, however, the 5-M33 system showed a left-shifted SASA distribution relative to the 20-Q33 system, indicating more effective shielding of the membrane surface by the branched architecture.

**Figure 10.**
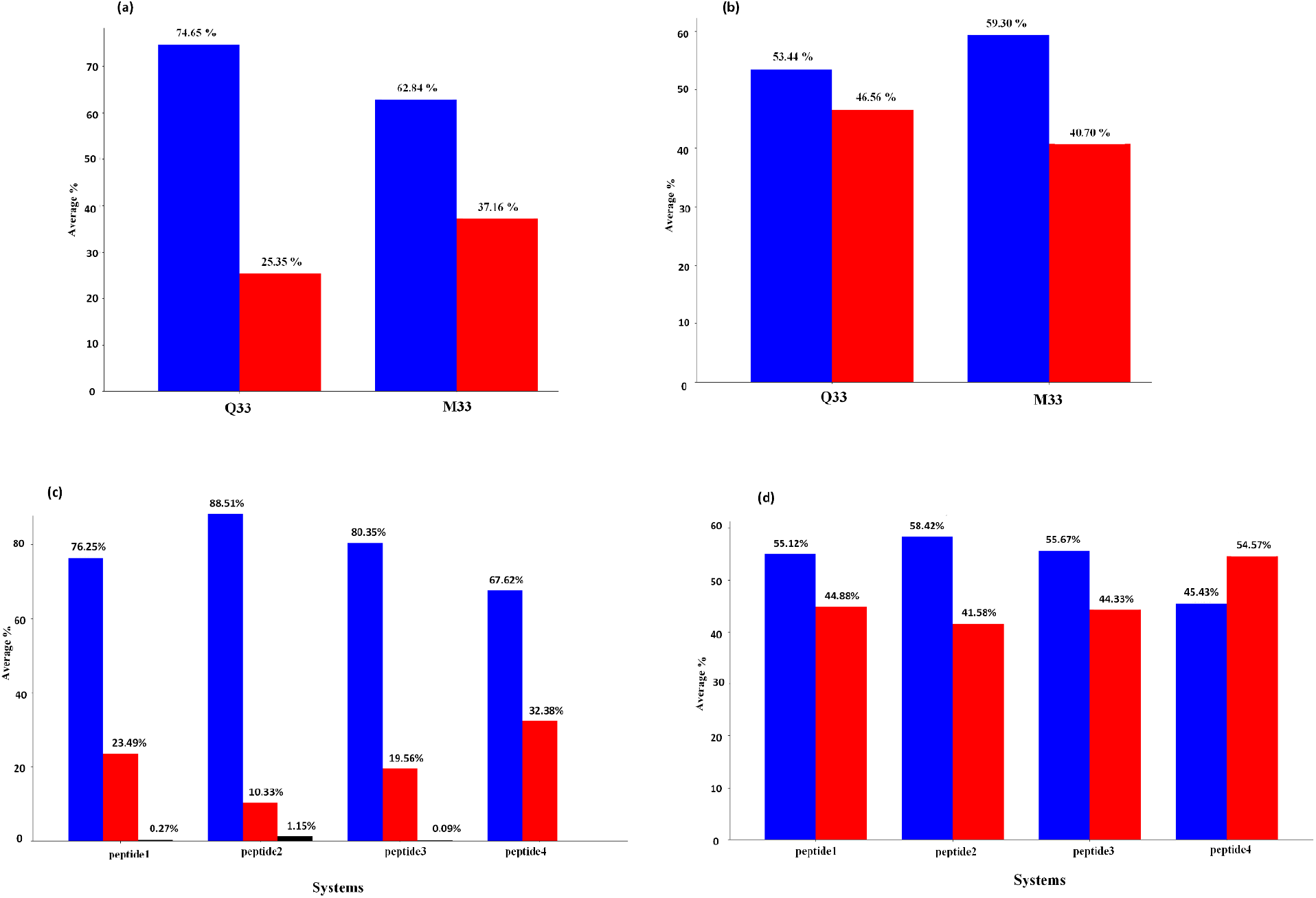
Secondary-structure evolution of Q33 and M33 in solution and membrane environments. Secondary structure was assigned with DSSP along the analyzed trajectories. Panels show representative profiles for Q33 and M33 in aqueous and membrane-associated systems. α-Helix, β-structure, turn/bend, and coil/disordered states are indicated according to the color legend. The secondary structure is color-coded as follows: red = α-helix, blue = disordered regions (loop, turn, bend), and black = β-sheet.

**Figure 11.**
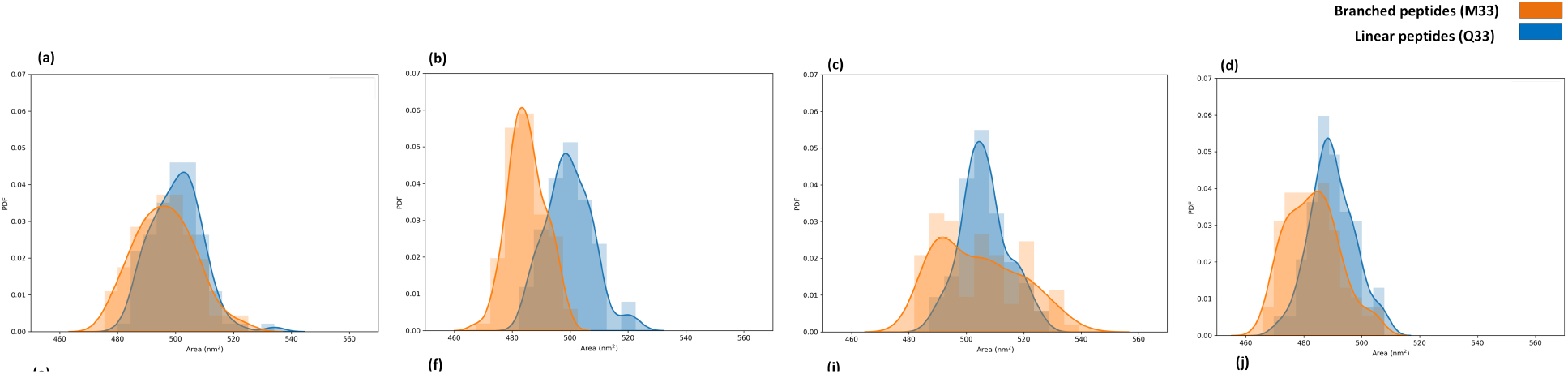
Membrane solvent-accessible surface area in Q33 and M33 systems. Probability density distributions of membrane SASA are shown for systems containing equivalent numbers of Q33 sequence units: (a) 4 Q33 and 1 M33, (b) 20 Q33 and 5 M33, (c) 32 Q33 and 8 M33, and (d) 40 Q33 and 10 M33. Lower SASA values correspond to greater shielding of the membrane surface by adsorbed peptides.

The broader SASA distributions observed for M33 suggest that membrane coverage by the branched constructs is more dynamic. This behavior may reflect fluctuations in the number of branches simultaneously engaged with the membrane, local reorientation of the scaffold, or transient hopping between neighboring lipid headgroups. In contrast, Q33 showed narrower SASA distributions, consistent with more localized peptide-rich clusters stabilized by peptide–peptide contacts.

Lateral peptide organization was further quantified using center-of-mass-based radial distribution functions (RDFs) and cumulative coordination numbers (CNs) calculated in the membrane plane (Table 2; Figures 12 and 13). For M33, the lowest-density system showed little short-range COM– COM enrichment and a main RDF peak around 3.8 nm, consistent with a relatively dispersed arrangement. Increasing the number of M33 molecules shifted the RDF toward shorter distances and increased first-neighbor coordination, but the overall CN values remained moderate. This indicates that M33 molecules become more laterally crowded at higher loading, while still avoiding the extensive close packing observed for Q33.

**Figure 12.**
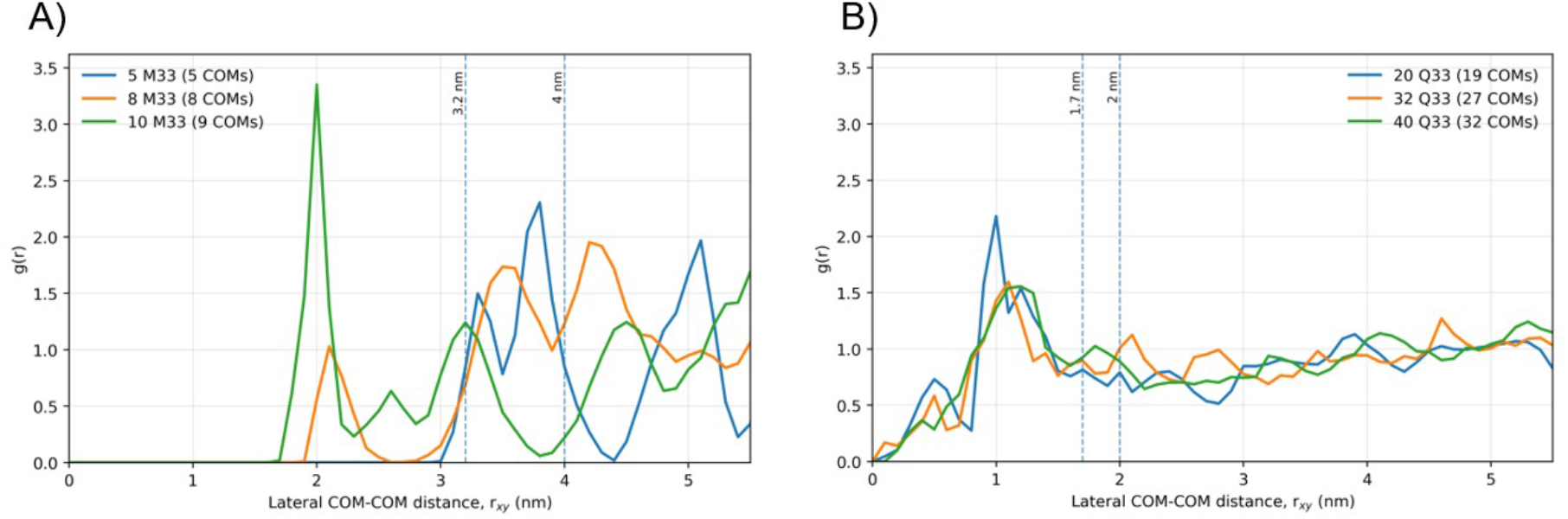
Lateral peptide–peptide radial distribution functions in the membrane plane. RDFs were calculated from peptide centers of mass for peptides associated with the same membrane leaflet. (A) RDFs for M33 systems. Dashed vertical lines indicate the M33 cutoffs used for close lateral association and first-neighbor coordination. (B) RDFs for Q33 systems. Dashed vertical lines indicate the corresponding Q33 cutoffs.

**Figure 13.**
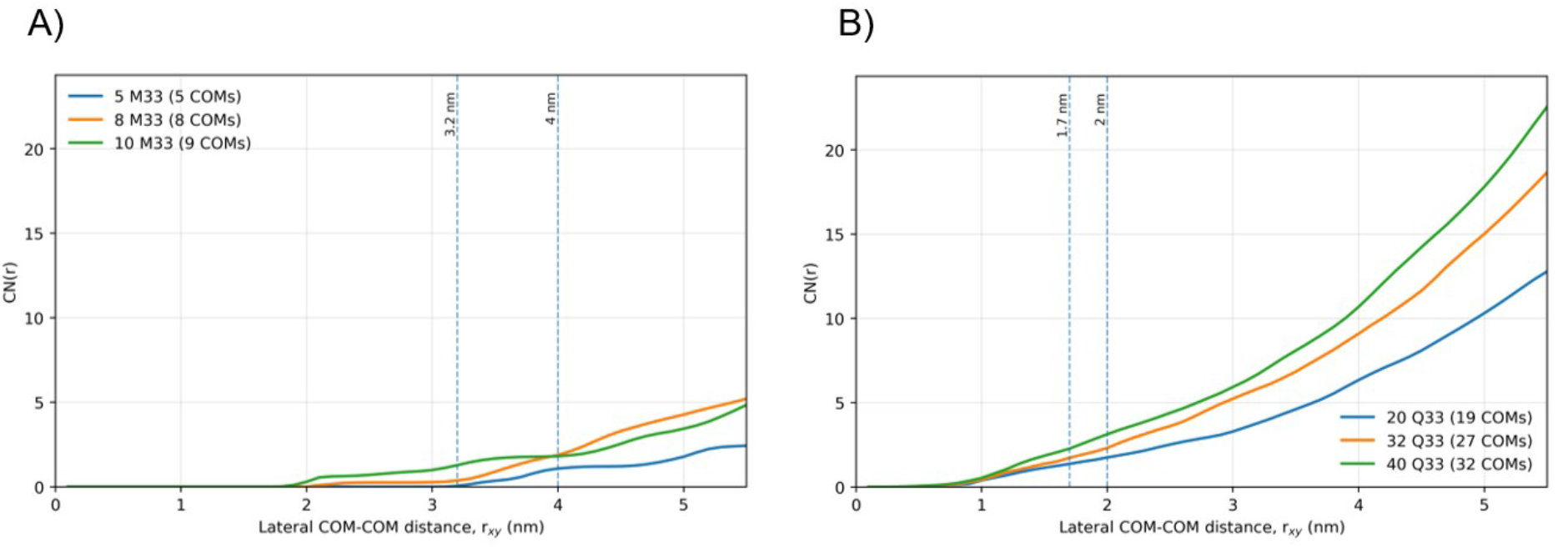
Cumulative coordination number analysis of peptide lateral organization. CN(r) reports the average number of neighboring peptides within a lateral center-of-mass distance r. (A) CN profiles for M33 systems. (B) CN profiles for Q33 systems. The analysis quantifies how local peptide crowding changes with peptide loading within each architecture.

**Table 2.** Lateral organization of membrane-associated peptides from center-of-mass radial distribution functions (RDFs) and cumulative coordination numbers.

| System | N | RDF peak r (nm) | RDF peak g (r) | Close-association CN | First-neighbor-shell CN |
| --- | --- | --- | --- | --- | --- |
| 5 M33 | 5 | 3.80 | 2.31 | CN(3.2 nm) = 0.05 | CN(4.0 nm) = 1.08 |
| 8 M33 | 8 | 4.20 | 1.95 | CN(3.2 nm) = 0.36 | CN(4.0 nm) = 1.88 |
| 10 M33 | 9 | 2.00 | 3.35 | CN(3.2 nm) = 1.28 | CN(4.0 nm) = 1.81 |
| 20 Q33 | 19 | 1.00 | 2.18 | CN(1.7 nm) = 1.37 | CN(2.0 nm) = 1.75 |
| 32 Q33 | 27 | 1.10 | 1.60 | CN(1.7 nm) = 1.73 | CN(2.0 nm) = 2.31 |
| 40 Q33 | 32 | 1.20 | 1.56 | CN(1.7 nm) = 2.27 | CN(2.0 nm) = 3.12 |

RDFs and cumulative coordination numbers were calculated in the membrane plane using peptide centers of mass for peptides associated with the same leaflet. Architecture-specific cutoffs were used to define close lateral association and first-neighbor shells: 3.2 and 4.0 nm for M33, and 1.7 and 2.0 nm for Q33.
<sup>N</sup> indicates the number of peptides included in the analysis on the selected leaflet.

Linear Q33 displayed a distinct lateral organization. The Q33 RDFs showed short-range maxima around 1.0–1.2 nm, and the cumulative coordination number increased progressively with peptide loading. These data indicate that Q33 peptides form compact peptide-rich regions on the membrane surface. Because Q33 systems contain many more independent peptide molecules than the corresponding M33 systems, absolute CN values should not be interpreted as a direct one-to-one comparison between architectures. Nevertheless, the architecture-specific RDF and CN profiles show that Q33 promotes short-range peptide crowding, whereas the larger tetra-branched M33 scaffold imposes greater spatial separation between membrane-associated constructs.

Peptide–membrane contact distance was assessed from the minimum distance between peptide heavy atoms and phosphate atoms of the upper membrane leaflet (Figure 14). Increasing peptide loading increased the probability of close peptide–membrane contact, reflecting more frequent engagement with lipid headgroups. When systems containing the same total number of Q33 sequence units were compared, Q33 and M33 showed similar minimum-distance distributions. Thus, peptide architecture strongly affects lateral organization and aggregation on the membrane, but it does not substantially alter the closest peptide–phosphate contact distance under the conditions studied.

**Figure 14.**
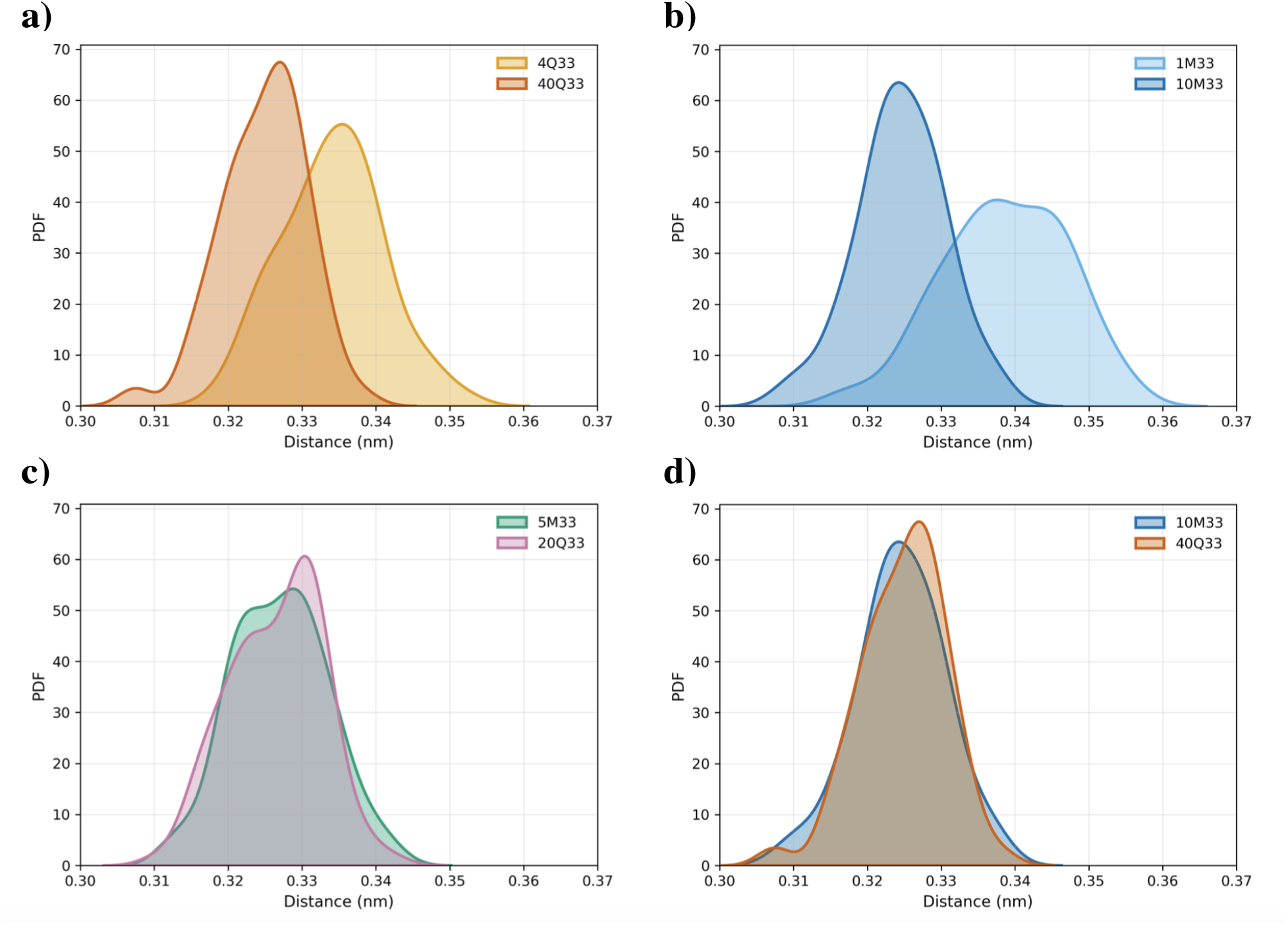
Peptide–membrane minimum-distance distributions. Minimum distances were calculated between peptide heavy atoms and phosphate atoms of the upper membrane leaflet. A 0.4 nm threshold was used to identify close peptide–membrane contact. The distributions report contact persistence or bound-state occupancy.

Overall, these analyses indicate that Q33 and M33 both bind efficiently to the anionic membrane surface, but they organize differently after adsorption. Linear Q33 favors compact, peptide-rich clusters, whereas M33 provides multivalent membrane engagement while maintaining larger intermolecular separation. This distinction helps explain how the branched scaffold can preserve strong membrane association while limiting uncontrolled self-aggregation.

### Membrane Binding Mode

Across the simulated systems, neither Q33 nor M33 produced persistent pore formation, complete peptide insertion, or peptide translocation through the bilayer within the sampled timescales. Instead, both peptide architectures remained predominantly associated with the membrane surface, maintaining close contact with lipid phosphate groups while preserving an interfacial rather than transmembrane orientation. In fact, the simulations capture an early membrane-association regime rather than the later stages of membrane disruption.

The observed binding mode is consistent with the initial phase of a carpet-like mechanism, in which cationic peptides accumulate on the membrane surface through electrostatic interactions with anionic lipid headgroups before large-scale bilayer destabilization occurs. In the present simulations, peptide adsorption was accompanied by local crowding and mild membrane deformation, but not by stable pore formation, membrane rupture, or extensive lipid extraction.

Therefore, the data support a surface-bound interaction model over the simulated timescales, while they do not exclude the possibility that membrane disruption could occur at longer times, higher peptide-to-lipid ratios, or in more compositionally complex bacterial envelope models.

The comparison between Q33 and M33 suggests that architecture affects how the surface-bound state is organized. Linear Q33 tends to accumulate as peptide-rich clusters, whereas branched M33 engages the membrane through a more spatially constrained, multivalent binding mode. Thus, both systems reach close membrane contact, but the lateral organization of the bound peptides differs substantially between the two architectures.

### Secondary-Structure Evolution

Secondary-structure analysis was performed to determine how peptide topology, peptide-peptide association, and membrane binding influence the conformational behavior of Q33 and M33. Experimental circular-dichroism and NMR studies have shown that Q33 can adopt α-helical structure in membrane-mimicking environments such as SDS micelles, whereas residue-level structural characterization of M33 is more challenging because of its branched architecture.

The initial models were constructed with α-helical Q33 segments, including the four branch segments of M33. Therefore, the simulations were used to assess whether this initial helicity was retained after relaxation in water or upon interaction with the membrane. In aqueous solution, both Q33 and M33 lost a substantial fraction of their initial helical content (Figure 10). Single Q33 retained α-helical structure only during the early part of the trajectory before converting to more disordered conformations, including turns, bends, and coil-like states. This behavior indicates that the α-helical state is not intrinsically stable for isolated Q33 in water on the simulated timescale.

Membrane association increased the persistence of Q33 helicity. In membrane-containing simulations, Q33 retained α-helical structure for a larger fraction of the trajectory, consistent with membrane-induced stabilization of an amphipathic helical conformation. Peptide-peptide contacts also contributed to conformational persistence in multi-Q33 systems. In particular, peptides involved in stable intermolecular contacts showed greater retention of ordered structure, suggesting that both membrane binding and local peptide packing can stabilize Q33 conformations.

M33 displayed a more heterogeneous conformational profile. Individual branches transiently retained helical segments, but the overall secondary structure evolved toward a mixture of helix, turn, bend, and coil states in both solution and membrane environments. Unlike Q33, M33 did not show a simple membrane-induced increase in helicity. Instead, its conformational organization appeared to be governed by the constraints of the branched scaffold, branch-branch electrostatic interactions, and the number and orientation of branches simultaneously engaging the membrane.

These results indicate that Q33 and M33 use different structural routes to membrane association. Q33 behaves as a short linear amphipathic peptide whose helicity is stabilized by the membrane and by peptide-peptide contacts. M33, in contrast, maintains membrane association despite a more heterogeneous secondary-structure ensemble, suggesting that persistent α-helicity is not required for its surface-bound state. For the branched peptide, topology, multivalent electrostatic binding, and controlled exposure of hydrophobic residues appear to be more important than a uniformly stable helical conformation.

Across the final 100 ns of the trajectories, the average α-helical content (mean ± SD) was 25 ± 0.19% for Q33 in solution and 42 ± 0.56 % for membrane-bound Q33, compared with 33 ± 0.46% and 34 ± 0.06% for M33, respectively.

### Residue-Level Hydration and Interfacial Orientation

Residue-level hydration was analyzed to identify which peptide regions remained solvent exposed and which became partially shielded from water upon membrane association (Figure 15). Hydration was quantified as the average number of water molecules within 0.4 nm of each residue over the analyzed trajectory window. Reduced hydration was interpreted as increased interfacial association or partial burial, whereas higher hydration indicated greater exposure to the aqueous phase. Because raw hydration values depend on residue size and chemical composition, the analysis was interpreted qualitatively and, where appropriate, compared across equivalent residues and environments rather than used as an absolute measure of insertion depth.

**Figure 15.**
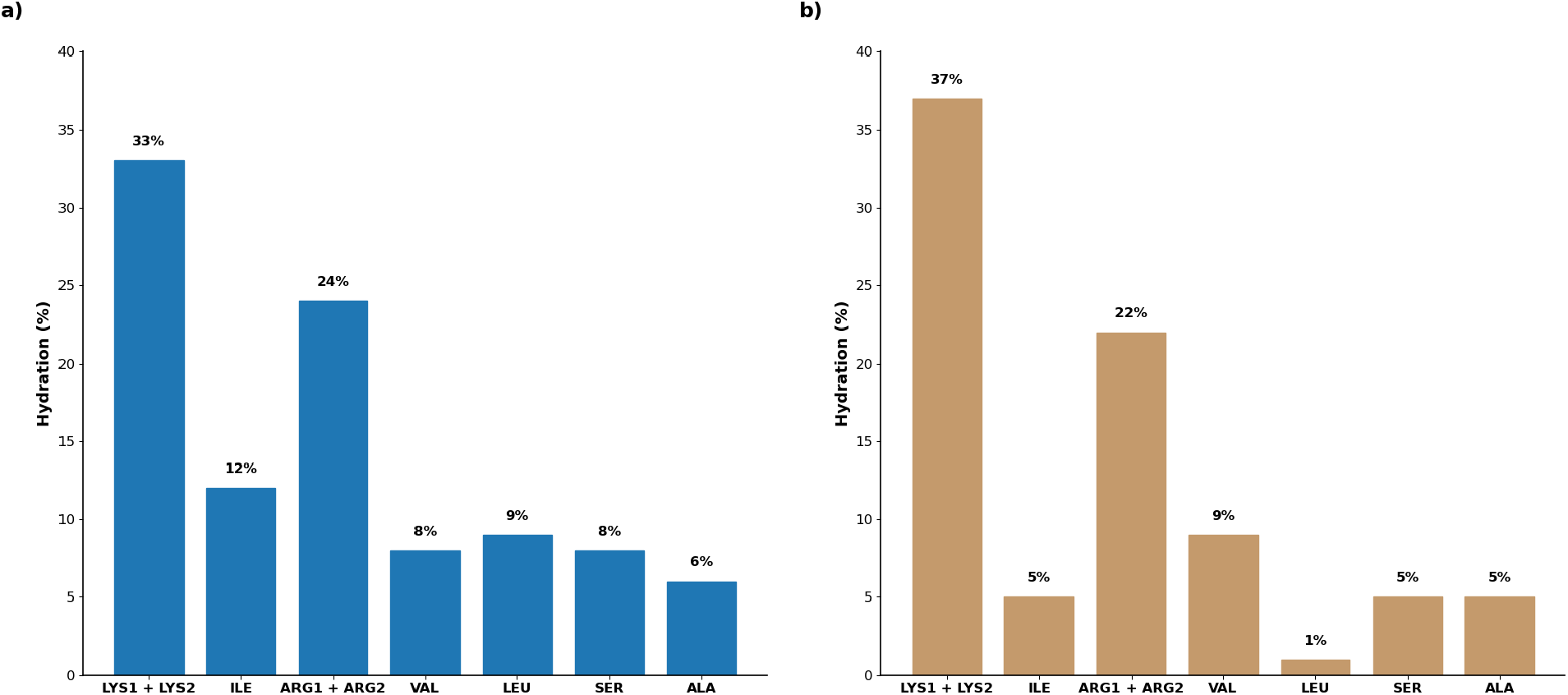
Residue-level hydration profiles of Q33 and M33. Hydration was quantified as the average number of water molecules within 0.4 nm of each residue over the analyzed trajectory window. Lower hydration indicates reduced solvent exposure, consistent with interfacial association or partial burial, whereas higher hydration indicates greater exposure to the aqueous phase. Values should be interpreted qualitatively because raw hydration depends on residue size and chemistry.

For Q33, the hydration profile was consistent with available experimental NMR attenuation data, which indicate preferential association of hydrophobic residues such as isoleucine and leucine with the micelle interface while charged residues remain more exposed to solvent ^26^.

In the simulations, hydrophobic residues, including isoleucine, leucine, and alanine, showed reduced hydration relative to the charged residues, consistent with closer association with the membrane interface or partial shielding by peptide-peptide contacts. In contrast, lysine and arginine retained higher hydration values, reflecting their continued exposure to water while participating in electrostatic interactions with anionic lipid headgroups.

For M33, for which equivalent residue-level attenuation data are not available, the simulations revealed a related pattern. Lysine and arginine residues displayed the highest hydration values, whereas alanine, serine, and valine showed lower hydration, suggesting preferential positioning closer to the lipid interface or within less solvent-exposed regions of the branched scaffold. This trend supports a model in which M33 maintains hydrated cationic branches near the aqueous phase and lipid headgroups while hydrophobic residues sample more interfacial environments.

These hydration results should be interpreted together with the contact, SASA, and lateral-organization analyses. In Q33, reduced hydration of hydrophobic residues may arise from both membrane association and peptide-peptide clustering. In M33, lower hydration of selected hydrophobic residues is consistent with membrane-facing branch orientations, but the branched topology makes the hydration pattern more heterogeneous across equivalent branches. Overall, residue-level hydration provides qualitative support for the proposed interfacial organization of both peptides and helps connect the simulations to experimental observations for Q33.

## CONCLUSIONS

All-atom molecular dynamics simulations show that linear Q33 and tetra-branched SET-M33 interact with an anionic membrane through related but distinct molecular pathways. Linear Q33 displays a greater propensity for peptide–peptide association, with extensive oligomerization observed at the membrane interface and under hydration-tuned sampling. These assemblies are stabilized by hydrophobic contacts together with backbone and charged-residue interactions, leading to localized peptide-rich regions on the membrane surface.

In contrast, the tetra-branched SET-M33 scaffold suppresses extensive self-aggregation while preserving strong membrane association. This behavior arises from the combination of multivalent electrostatic attraction to anionic lipid headgroups, steric and topological constraints imposed by the branched core, and reduced exposure of hydrophobic/core regions to neighboring peptide molecules. Stable SET-M33–SET-M33 contacts were observed only under specific orientations, whereas widespread oligomerization comparable to that of Q33 was not detected.

Both peptide architectures remained predominantly surface associated over the simulated timescales, with no persistent pore formation, complete insertion, or translocation through the bilayer. Secondary-structure and hydration analyses further indicate that Q33 is more strongly stabilized by membrane-induced helicity and peptide-peptide contacts, whereas SET-M33 maintains membrane association through a more heterogeneous, topology-dependent conformational ensemble.

Overall, these results suggest that branched topology reduces uncontrolled peptide self-aggregation while maintaining efficient membrane engagement. This provides an atomistic rationale for the distinct behavior of SET-M33 relative to its linear branch sequence and supports branching as a useful design strategy for membrane-active antimicrobial peptide constructs.

## Supporting Information

Representative snapshots of Q33 peptide aggregation at the membrane.

## Corresponding Author

* Walter Rocchia

## Author Contributions

S.S.: Conceptualization, methodology, validation, formal analysis, investigation, data curation, visualization, writing-original draft, and writing-review and editing. M.R.: Methodology, formal analysis, investigation, visualization, and writing-review and editing. S.F.: conceptualization and supervision. C.F.: supervision and review. A.P.: supervision. W.R.: conceptualization, supervision, project administration, funding acquisition, and writing-review and editing. The manuscript was written through contributions of all authors. All authors have given approval to the final version of the manuscript.

## ACKNOWLEDGMENTS

The financial support received for three years doctoral research by the European Union Horizon Europe MSCA DN-ID SSBB doctoral network. This work was supported by grant number 101073263. doi: https://cordis.europa.eu/project/id/101073263. We also would like to acknowledge the access to the CINECA supercomputers, thanks to EUROHPC and ISCRA initiatives.

## ABBREVIATIONS

CN: coordination number
COM: center of mass
DOPE: 1,2-dioleoyl-sn-glycero-3-phosphoethanolamine
DPPE: 1,2-dipalmitoyl-sn-glycero-3-phosphoethanolamine
DPPG: 1,2-dipalmitoyl-sn-glycero-3-phospho-(1′-rac-glycerol)
DSSP: Define Secondary Structure of Proteins
LPS: lipopolysaccharide
MAP: multiple antigen peptide
MD: molecular dynamics
NPT: constant number of particles, pressure and temperature ensemble
NVT: constant number of particles, volume and temperature ensemble
PBC: periodic boundary conditions
PDB: Protein Data Bank
PME: particle mesh Ewald
RDF: radial distribution function
RESP: restrained electrostatic potential
RMSD: root-mean-square deviation
RMSF: root-mean-square fluctuation
SASA: solvent-accessible surface area
TIP3P: transferable intermolecular potential with three points
VMD: Visual Molecular Dynamics

---

**Figure S1.**
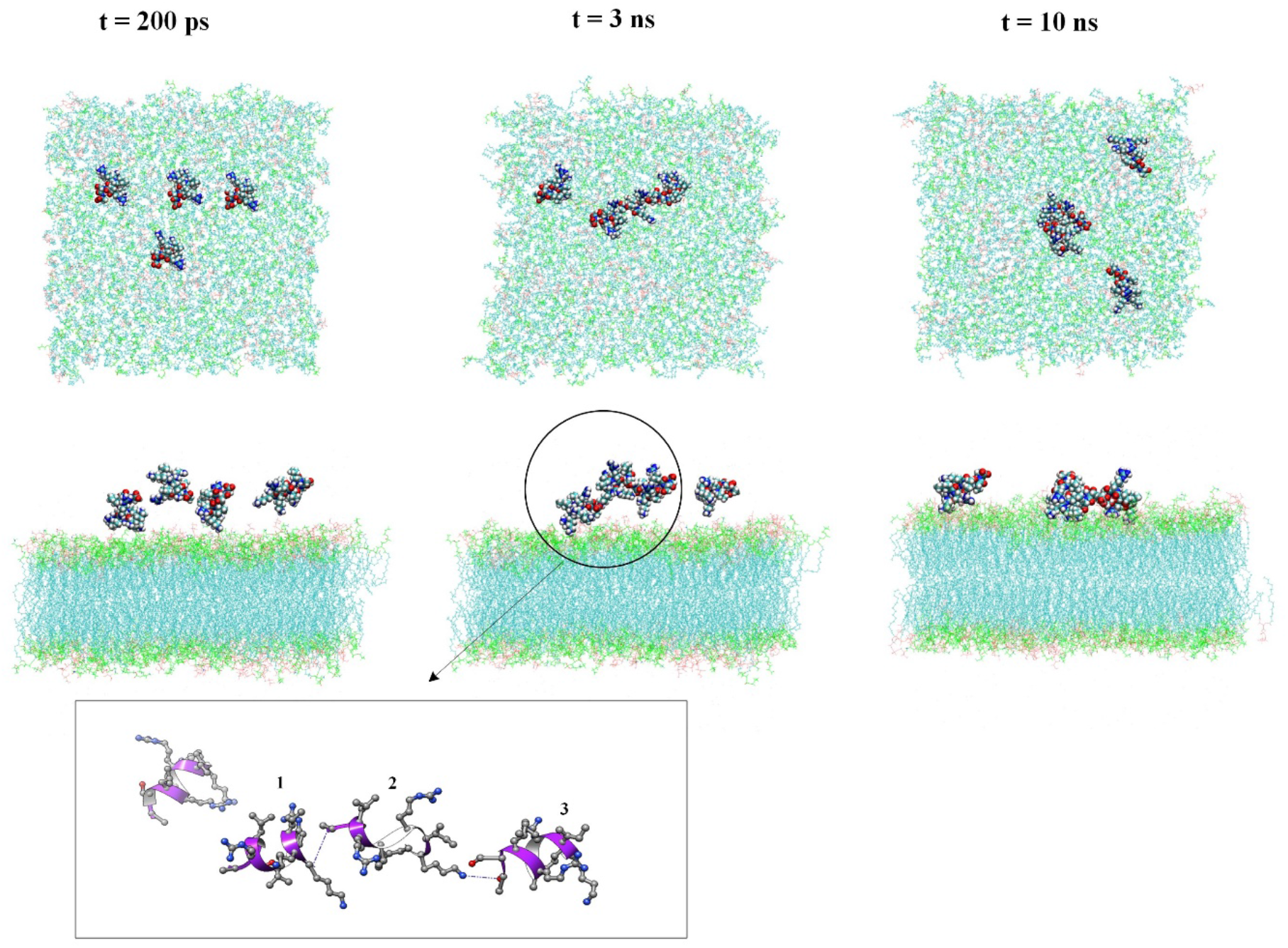
Representative snapshots of one replica of MD simulation for the system of 4-Q33-membrane at 0.2, 3, and 10 ns after removal of positional restraints. The system contains four Q33 peptides initially positioned above the membrane. Within 3 ns, three peptides aggregate and adsorb on the bilayer surface.

**Figure S2.**
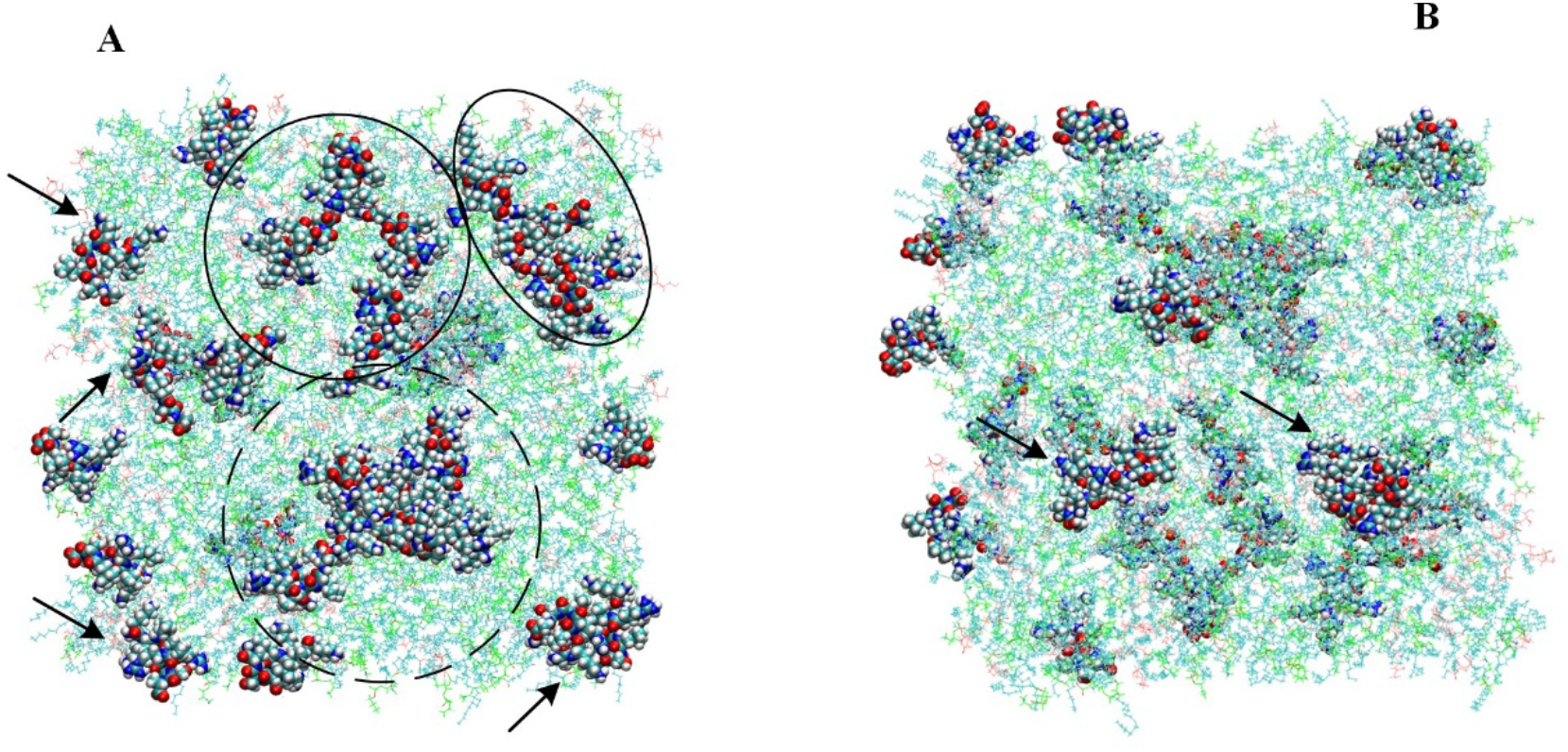
Representative snapshots of the Q33-membrane system containing 32 Q33 peptides. After 300 ns of simulation, peptides interact with both leaflets of the bilayer, forming dimers (arrows), tetramers (circles), and octamers (dashed circle). Panels A and B correspond to the upper and lower membrane leaflets, respectively.

**Figure S3.**
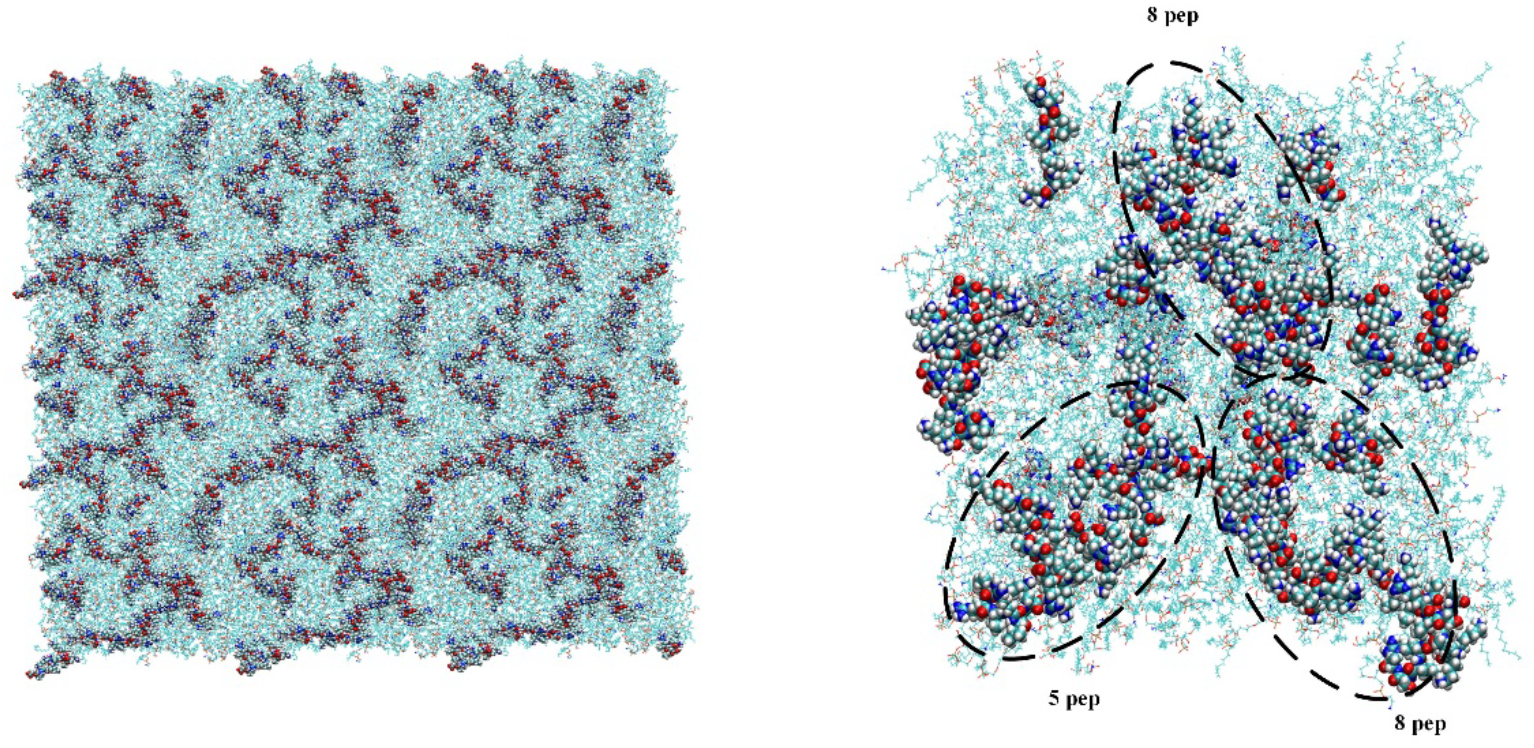
Representative snapshots of the Q33-membrane system containing 40 Q33 peptides. After 300 ns of simulation, peptides form octamers and pentamers on the membrane surface.

